# Single-trial-based Temporal Principal Component Analysis on Extracting Event-related Potentials of Interest for an Individual Subject

**DOI:** 10.1101/2021.03.10.434892

**Authors:** Guanghui Zhang, Xueyan Li, Yingzhi Lu, Timo Tiihonen, Zheng Chang, Fengyu Cong

## Abstract

Temporal principal component analysis (t-PCA) has been widely used to extract event-related potentials (ERPs) at the group level of multiple subjects’ ERP data. The t-PCA is, however, poorly employed to isolate ERPs from single-trial data of an individual subject. Additionally, the effects of varied trial numbers on the yields of t-PCA have not been systematically examined. To fill both gaps, in an emotional experiment (22 subjects), we use t-PCA and Promax rotation to extract interesting P2/N2 from single-trial data of each subject with consecutive increasingly trial numbers (from 10 to 42) and all trials, respectively. Besides, time-domain analysis and other two group t-PCA strategies (trial-averaged and single-trial) are also employed to isolate ERPs of interest from all subjects. The results indicate that the proposed technique produces the internal consistent measure of N2 from few trials (i.e., 19) as from all trials compared with the other three approaches (more than 30 trials). As for P2, all approaches yield internal-subject consistent effect after approximately 33 trials are included in the average, but Cronbach’s alpha values for the proposed technique are higher than the other two group PCA strategies over varied trials. Combined, the yields provide evidence that the proposed approach may efficiently temporally filter the data to extract more reliable and stable ERPs for an individual subject.

## 1 Introduction

Event-related potentials (ERPs) have been widely used to investigate brain cognitive processes in many fields, such as language, neuroscience, physiology, psychology, and so forth (Luck, 2014). Traditionally, ERPs are obtained by averaging tens or hundreds of single-trial EEG data under same experimental condition, improving signal-to noise ratio (SNR) of ERPs (Luck, 2014; Handy, 2005). SNR of ERPs is proportiona to the square root of trial numbers N, i.e., 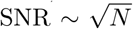 (Luck, 2014). More trials not only insignificantly improve SNR, but also making participants feel more fatigued that has a side effect on the task execution and enhances other signals of non-interest (e.g., alpha band). Therefore, it is critical and necessary to keep balance between data quality and experimental time by optimizing the trial numbers involved in experiments.

Many previous studies have gave data-driven evidence and guideline on how many trial numbers should be contained in some frequently used ERP experimental paradigms. Several typical ERP components have been investigated in these studies, for example, N2, P3, N1, P1, P2, error-related negativity (ERN), Late positive potential (LPP), N170, and error positivity (Pe) (Clayson, 2020; Larson et al., 2010; Huffmeijer et al., 2014; Pontifex et al., 2010; Olvet and Hajcak, 2009; Fischer et al., 2017; Thigpen et al., 2017; Cohen and Polich, 1997; Rietdijk et al., 2014; Kleene et al., 2021; Clayson and Larson, 2013). Most of these studies have quantified how to use minimum trial numbers to obtain the stable and reliable measures of ERPs of interest based on their original waveforms, etc. By contrast, the examination of the trial numbers influence on the decomposition of temporal principal component analysis (t-PCA) is poorly reported in past studies.

Regarding t-PCA application, two strategies are frequently used to extract ERPs of interest from all subjects’ datasets at the group level. In the first strategy, the pre-processed single-trial EEG data are initially averaged under the same condition and then the averaged datasets of all subjects are concatenated across channels, conditions, and subjects to form a matrix. The size of the matrix is time samples multiply by the combinations of electrodes, conditions, and subjects. Afterward, t-PCA and rotation method are performed on the matrix to extract factors ^1^ associated with ERPs of interest. In other words, this strategy is group t-PCA analysis for the averaged ERP datasets (Fogarty et al., 2020; Male and Gouldthorp, 2020; Bonmassar et al., 2020; Dien, 2012, 1998; Dien et al., 2007; Kayser and Tenke, 2003, 2006a,b). In the second strategy, ERPs of interest are extrcted from the matrix formed by single-trial EEG datasets of all subjects under all experimental conditions. Here, time samples are variables, the combinations of channels, conditions, trials, and subjects are observations. The related decomposition is named single-trial t-PCA analysis at the group level (Rushby and Barry, 2009; MacDonald and Barry, 2017; MacDonald et al., 2015; Rushby et al., 2005).

Noticeably, two things need to notice for both group t-PCA strategies. On the one hand, the core ideas of the two strategies are high similar that ERPs of interest are extracted from datasets of all subjects. If all the factors decomposed from the same ERP are projected to electrode fields (i.e., in microvolts) and single-trial EEG data are averaged before or after PCA decomposition, we might obtain a significant similar result of both strategies because PCA is a linear decomposition (Wold et al., 1987; Comon and Jutten, 2010). In the above, a certain estimated number of sources should also be used for both strategies. The further comparison and discussion between the two group t-PCA strategies can be found in the results. On the other hand, both group t-PCA strategies follow the same assumption (i.e., blind source separation, BSS) (Cong et al., 2011a,b; Makeig et al., 1997, 1999): **Z** = **a**_*m*,1_·**s**_1_(*t*) + … + **a**_*m,r*_·**s**_*r*_(*t*) +…+ **a**_*m,R*_·**s**_*R*_(*t*) = **AS. Z** is a spatially concatenated matrix from either trial-averaged or single-trial datasets of all subjects under different conditions; the rows of **Z** are time samples and the multiplications of the others (e.g., subjects, channles, conditions, etc.) are in columns. *s*_*r*_(*t*) represents stimulus-locked, spontaneous, or noise sources. *a*_*m,r*_ is coefficient between *m*^*th*^ electrode and *r*^*th*^ source. We consider that the sources are invariant over all subejcts, that is, **S**_*r*_ = **S**(1)_*r*_ = … = **S**(*p*)_*r*_ = … = **S**(*P*)_*r*_ (r is the source sequence and P is the number of subjects). However, not all the sources in brain cortex for different subjects are the same even they respond to the same stimuli under the same environment because brain functions among people are differ a bit. This gives us motivation to explore the intersting ERPs from single subject’s EEG data.

In order to explore P2/N2 from single-trial EEG data of individual subject and investigate the effect of the increasing trials on t-PCA yields in an amotional ERP experiment, the following steps are involved (see Fig. 1). Firstly, the datasets of consecutive increasingly trials (i.e., from 10 to 40) are stacked along electrodes (i.e., spatial) of different conditions to form a matrix for singletrial EEG of an individual subject. Secondly, t-PCA and Promax rotation are employed to decompose the matrices separately. Next, the factors associated with ERPs of interest for each subject are chosen and projected to the electrode fields (i.e., in microvolts). After that, the back projections of single trials are averaged separately under each experimental condition for each subject. Finally, the mean amplitudes of the desired ERPs are quantified within a predefined time-window at some electrodes. Meanwhile, we also separately calculate the similarities of topographies between different subjects for varied trial numbers to evalute the internal consistent of P2/N2. The statistical results for increased trials and related similarities are compared with these of all trials. Besides, we also measure the correlation cofficients (CCs) of P2/N2 between the averages of smaller trials (from 10 to 42) and the grand average (all trials), and compute the Cronbach’s alpha for the increased trials.

**Fig. 1.**
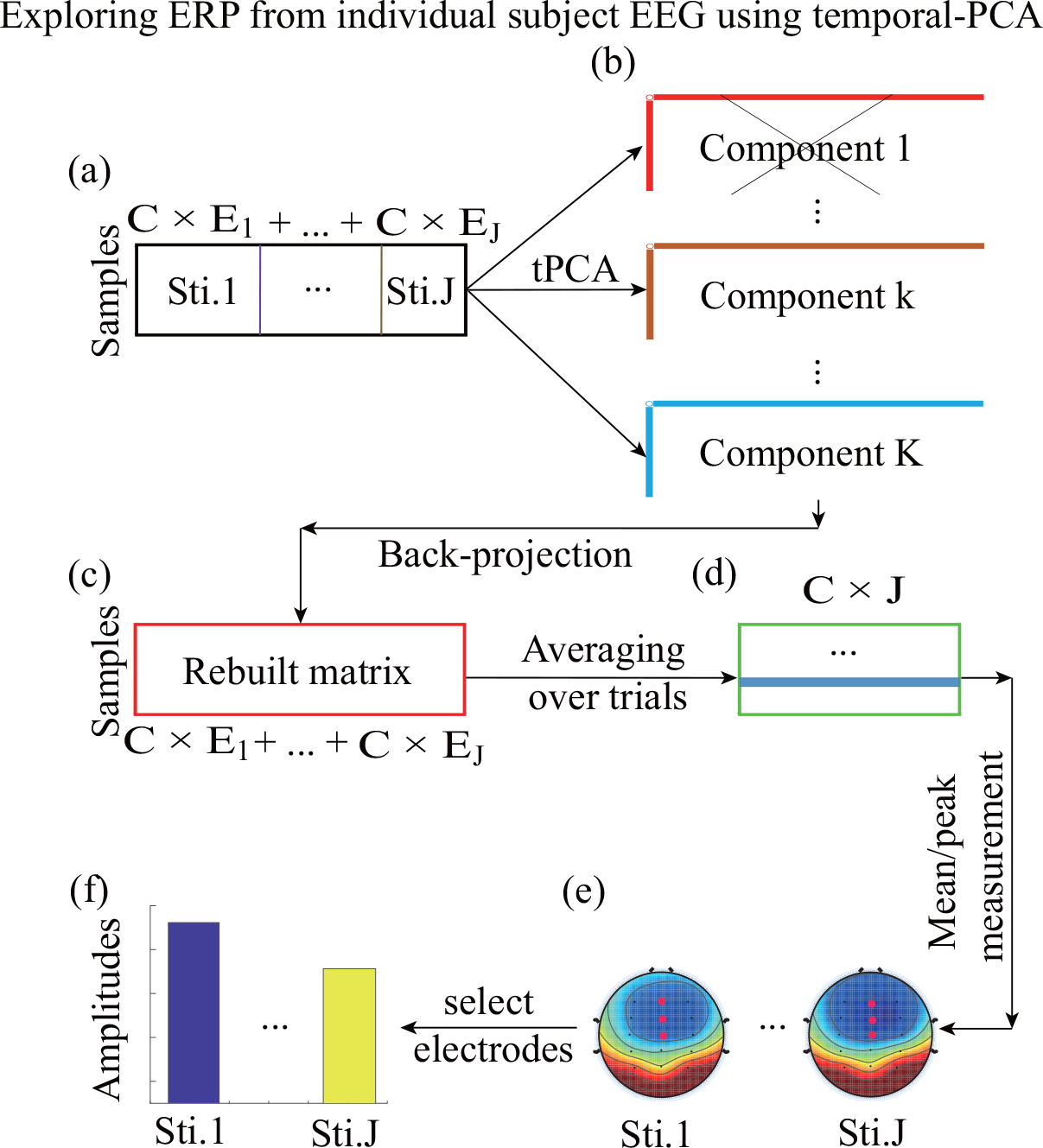
The illustration of the proposed technique. (a) The single-trial EEG for individual subject are arranged into a matrix over channels (trial numbers of conditions 1, …, and *J* are *E*_1_, …, and *E*_*J*_, respectively; *C* is number of electrodes). (b) The matrix are decomposed into sums of R components by using temporal-PCA and Promax rotation. (c) Projecting the selected components back to electrode fields for building a new matrix. (d) Reconstructed waveforms of single-trial EEG are averaged under each condition. (e)-(f) Calculating the amplitudes of the desired ERP for different conditions within a defined time-window at some specific electrodes.

Furthermore, conventional time-domain analysis and two group PCA strategies as mentioned-above are considered as comparison techniques, and they are also applied to the analysis of the same pre-processed EEG datasets for increasing trials and all trials. The pre-processed EEG datasets and codes used in this study are available from: http://zhangg.net/publications/.

## 2 Data description and method

In this section, we first briefly describe the ERP experiment, which are reanalyzed here and has been published (Lu et al., 2016). Afterward, the model and procedures of PCA are introduced.

### 2.1 Paticipants

A total of 22 healthy undergraduate students come from Shanghai University of Sport participate in the study as paid volunteers. This experiment is approved by the local ethics committee and all participants sign their written informed consent before the experiment. The age of paticipants is 22.05 ± 1.53 years old (range: 20 to 24), including 10 males and 12 females.

### 2.2 Task

Participants are required to respond to different emotional pictures in a modified oddball distinction task. The task in each trial starts with a black fixation (fixed time is 300 ms) in the center of the white computer screen. In the following, a blank white screen is displayed with a random time-window between 500 ms and 1500ms. Afterward, the stimuli are displayed to participants. Stimuli will be disappeared after 1000ms or terminated by participants who press a key (‘J’ for the deviant pictures and ‘F’ for standard ones). Another blank screen with 1000ms will follow the response before start next trial.

Six blocks are included in the experiment. 100 trials are contained in each block that comprises of 70 standard stimuli and 30 deviant stimuli. A natural scene of a chair is used for standard stimulus. Deviant stimuli include 10 moderately/extremely disgusting pictures, 10 moderately/extremely fearful pictures, and 10 neutral pictures, separately.

### 2.3 EEG recording and preprocessing

Nineteen electrodes (F3, FC3, C3, CP3, P3, Fz, FCz, Cz, CPz, Pz, F4, FC4, C4, CP4, P4, TP9, TP10, VEOG, and HEOG) are used to record EEG based on the international 10-20 system with 1000 Hz sampling rate. EEG recordings are referenced at FCz (Brain Products GmbH, Germany). All impedances are less than 5 k*Ω* for each sensor of each subject.

The collected EEG data are pre-processed offline. Firstly, the left and right mastoids are set to offline references, and the sampling rate is set to 500Hz. Secondly, EEG data are filtered by an infinite impulse response (IIR) band-pass filter: lower cut-off is 0.1 Hz, higher cut-off is 30 Hz, and slope: 48 dB/oct. Next, the filtered EEG data are segmented from 200 ms before stimulus onset to 900 ms after stimulus onset. Those epochs’ datasets whose magnitudes exceeded ±80 *µ*V are discarded and the remaining epochs are baseline corrected. Finally, EEG data of all epochs are filtered by wavelet filter (Cong et al., 2015; Zhang et al., 2020) to improve signal-to-noise ratio (SNR). The parameters are set for wavelet filter as below: the number of decomposition level iss 10; the select mother wavelet is ‘rbio6.8’; the detail coefficients at levels 5, 6, 7, 8, and 9 are used for signal reconstruction. The preprocessed EEG data of 20 subjects are involved in analyzing N2 and 17 subjects for P2. It should be noted that two stimuli for the neutral condition (i.e., neural disgusting and neural fearful) are merged to one in previous study (Lu et al., 2016).

The details for data collection and experiment can be found in (Lu et al., 2016).

### 2.4 Mathematical model and procedures for principal component analysis

In this subsection, we introduce the model for PCA and procedures for the application of PCA to ERP/EEG datasets.

#### 2.4.1 Mathematical model for principal component analysis

The application of PCA to a spatial-stacked matrix **Z** (the rows are time points and the channels from each condition/subject/trial (optional) are in columns) obtained from EEG/ERP dataset of either single subject or multiple subjects can be illustrated (Zhang et al., 2020; Cong et al., 2011a,b; Makeig et al., 1997, 1999):

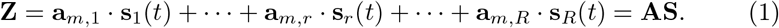

In Eq. 1, the number of observed signals is larger than that of sources, i.e., it is an over-determined model, and we usually change this over-determined model to determined model by some methods, for example, the accumulative explained variance (Cong et al., 2011a,b; Zhang et al., 2020).

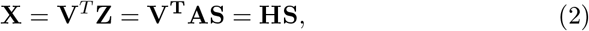

where **V**^*T*^ is the dimensionality reduction matrix obtained by applying some PCA algorithms to matrix **Z**^*T*^ and **H** is a mixing matrix.

In applications of PCA, we seek an un-mixing matrix **W** by using some rotation algorithms, such as Promax, Varimax rotations, and so forth (Richman, 1986; Hendrickson and White, 1964; Kaiser, 1958). Once the un-mixing matrix is generated, its inverse matrix **B** = **W**^−1^ is used to estimate **H**. And we can also use the unmixing matrix **W** to convert **X** into an estimated component matrix:

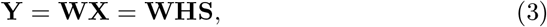

here, each row of the estimate matrix **Y** represents topography of each source (i.e., rotated factor scores); **C** = **WH** is a global matrix.

Generally, we need to choose several factors derived from **X** for further analysis (Comon and Jutten, 2010), and thus, the theory of back-projection is used to analyze these factors simultaneously as applied in the previous studies (Dien, 1998; Cong et al., 2011a,b; Makeig et al., 1997, 1999). In the matrix-vector form, the back-projection is the outer product of *k*^*th*^ column of **B** with *k*^*th*^ row of estimated factor matrix **Y**:

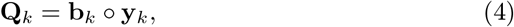

where **Q**_*k*_ represents the back-projected signals at all the electrodes for *k*^*th*^ selected factor; ‘o’ denotes the outer product of two vectors.

Under global optimization, only one nonzero element exists in each row and each column of matrix **C** so that the extracted *k*^*th*^ factor can uniquely represent *j*^*th*^ unknown scaled source (Cong et al., 2011a,b):

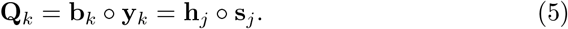

Regarding the original over-determined model in Eq. 1, *k*^*th*^ factor generated from **Z** is selected to project back to electrode fields and this procedure can be described as below:

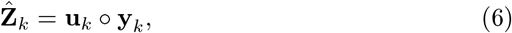

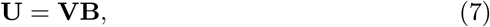

where **U** is the estimation of **A** and its each column demotes time course for *k*^*th*^ factor (i.e., the *k*^*th*^ rotated factor loading).

In the application of PCA, a desired ERP is often decomposed into several factors because the latencies of the ERP vary among different subjects. Therefore, those factors need to be projected back to the electrode fields simultaneously based on the following rule (Cong et al., 2011a,b; Zhang et al., 2020):

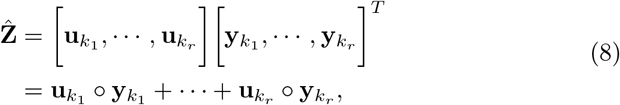

where *k*_1_, …, and *k*_*r*_ (1 ≤ *k*_*r*_ *< R*) represent the orders of the identified factors.

#### 2.4.2 Principal component analysis procedures

To better extract ERPs of interest from preprocessed ERP/EEG datasets by using PCA and Promax rotation for further analysis, the following steps are involved: arrange pre-processed ERP/EEG datasets into a matrix, estimate the number of sources, select an optimal rotation method, identify factors of interest, and project the identified factors to electrode fields (i.e., in microvolts). Here, trial-averaged data are taken as an example to explain the PCA procedure on the extractions of ERPs.

The first step is to arrange ERP datasets into a matrix. The recorded EEG datasets are born with space and time dimensions, and thus, for multicondition and multi-subject trial-averaged datasets, two types of matrices can be organized (Dien, 2012; Dien et al., 2007, 2005). For the first type of matrix, it is formed by concatenating multi-condition and multi-subject datasets over electrodes, that is, time points are variables and the combinations of electrodes and subjects-conditions are observations. The related PCA procedure is named as ‘t-PCA’. For the second type of matrix, electrodes are variables and the other variances (i.e., time points and subjects-conditions) are merged as observations. Likewise, we call the performance of PCA on this type of matrix as ‘s-PCA’. In this study, the former type is formed based on the following reasons. For one thing, regarding the performance of t-PCA and s-PCA, t-PCA can yield overall better results than s-PCA (Dien, 1998, 2012). For another thing, the desired ERP is easily mixed with others in the spatial domain to some degree due to the volume conduction (Dien, 2012).

The second step is to estimate the number of sources. The purpose of this step is to transform the over-determined model in Eq. 1 into the determined model in Eq. 2. Several approaches have been developed to realize this goal, such as Parallel Test (Horn, 1965; Dien, 2010a), gap measure (He et al., 2010; Cong et al., 2013), and cumulative explained variance (Zhang et al., 2020; Huster and Raud, 2018; Arbel et al., 2013). Here, we use the last approach to estimate the number of sources and this approach is to calculate the percentage between the sums of first *R* lambda values (one lambda corresponds to one factor) over the sums of all lambda values:

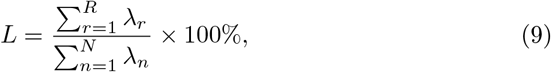

where *R* is the estimated source number; *N* is the number of variables/time points here (*N* ≥ *R*); The lambda values are in descending order: *λ*_1_ ≥ … ≥ *λ*_*n*_ = … = *λ*_*N*_ = *σ*^2^. Once *L* is given with a specific value, for example, 95%, 99%, and so on, the corresponded number of sources can be then obtained.

The third step is to select an optimal rotation method. The goal of rotation is to rearrange the factors into simple and interpretable structures so that one factor corresponds to one ERP (Dien, 2012). Some rotation methods, such as Promax, Varimax, and Infomax, can be utilized. The results of actual and stimulated ERP datasets indicated that Promax rotation showed more accurate results than Varimax rotation (Dien et al., 2005; Dien, 1998; Dien et al., 2003). And, simulation ERP studies revealed that Promax generated overall better results for t-PCA, and Infomax yields better separation for sPCA (Dien et al., 2007; Dien, 2010b).

The fourth step is to identify factors of interest. Although some pre-processing steps are utilized to improve SNR of ERP/EEG signals, that is, some artefacts are removed, like DC offset, slow drift of sensors, eye movement, line noise, and muscle contraction (Jung et al., 2000; Delorme et al., 2007; de Cheveigné and Nelken, 2019; de Cheveigné, 2020; Sai et al., 2017), the preprocessed data still contain spontaneous EEG brain activities, ERP components of non-interest, ERP components of interest, and noise activities. Therefore, we need to identify those factors associated with ERPs of interest for further analysis. Generally, the identification of desired factors is based on the two aspects (Dien, 2012; Barry et al., 2020; Zhang et al., 2020): (1) the polarity and latency of temporal factor; (2) the polarity and topographical distribution of spatial factor.

The fifth step is to project the identified factors back to the electrode fields (i.e., rescale them to microvolts). When performing PCA on an ERP dataset of multiple subjects, an ERP of interest may be decomposed into several factors because the differences are found in latencies or phases of this ERP across different subjects to some degree. Therefore, all of the decomposed factors related to this ERP of interest should be back-projected onto electrode fields (Dien, 2012, 1998; Zhang et al., 2020).

## 3 Data analysis

In the current study, ERPs of interest, i.e., P2 and N2, are reanalyzed, which have been reported in (Lu et al., 2016). For the proposed technique and other three alternative approaches, the two ERPs are separately quantified from all trials and consecutive increasing trials (i.e., 10, 11, …, 41, and 42 trials). Notably, 10 is applied to ensure the number of observations is larger than that of variables when using PCA. 42 is used to the upper number because it is the minimum trial number for different subjects under all conditions.

In the following, we detail how to use the four techniques to extract P2/N2 from either all trials or increased trial numbers.

### 3.1 Method 1 (‘M1’): Proposed technique

In order to better identify the factor(s) extracted from pre-processed EEG dataset of single subject by utilizing t-PCA and Promax rotation, a timewindow is set for P2 (130-190 ms) and N2 (190-310 ms) separately.

The following steps are taken when using proposed approach to extract P2/N2 from the pre-processed EEG dataset for each subject.

1. Single-trial EEG dataset of *i*^*th*^ subject is arranged into a matrix **Z**^(*i*)^ with size of *T ×* (*C ×* (*E*_*i*,1_ + …+ *E*_*i,J*_)), *T* is the number of time points, *C* is the number of electrodes, and *E*_*i,j*_ is the number of trials for *j*^*th*^ condition (*j* = 1, …, *J* − 1, and *J*). Noticeably, for the analysis of all trials, *E*_*i*,1_, …, *E*_*i,J*−1_, and *E*_*i,J*_ may be variant. As for the analysis of adding trial numbers, they are the same that are equal to 10, 11, …, 41, and 42, separately.
2. *T* factors are extracted by performing PCA on the matrix (**Z**^(*i*)^)^*T*^ and then *R*^(*i*)^ factors are retained and rotated in Matlab environment (Version 2018b, the Mathworks, Inc., Natick, MA; functions: *pca.m* and *rotatefactors.m*; *R*^(*i*)^ ≤ *T*); Here, aiming at successfully separating ERPs of interest with others, the number of retained factors is set to 40, which account for more than 99%, in both all trials and increased trials.
3. These factors associated with P2/N2 are identified for the next procedure according to following aspects. The first thing, the latency of *r*^*th*^ temporal factor 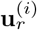 fell in the predefined time window. The other one, topographic distribution of *r*^*th*^ spatial factor 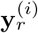 is in accordance with that of P2/N2.
4. The identified factors are projected onto electrode fields for correcting their variance and polarity indeterminacies and reconstruct a new matrix 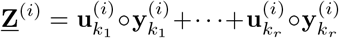. Single-trial EEG data are separately averaged for each experimental condition.
5. Repeating steps (1) - (4) until the single-trial EEG datasets of all subjects are decomposed.

### 3.2 Method 2 (‘M2’): Conventional time-domain analysis

For the all trials analysis, single-trial EEG data for each condition are averaged to obtain the waveform of ERPs. Likewise, for the increased trials analysis, EEG data are also averaged for different conditions when the trial number is 10, 11, …, 41, and 42, separately.

### 3.3 Method 3 (‘M3’): Group PCA for trial-averaged ERP data

We use the same procedure in ‘M1’ to quantify P2/N2 in this method but a bit changes in steps 1 and 4. Specifically, in the step 1, the trial-averaged ERP data for different subjects under different experimental conditions are organized into a big-size matrix. The rows of **Z** are time points, and columns are the multiplications of electrodes, conditions, and subjects. In the fourth step, we only project the selected factors onto electrode fields to reconstruct the waveforms of P2 or N2.

### 3.4 Method 4 (‘M4’): Group PCA for single-trial EEG data

The key discrimination between this method and M1 is that single-trial EEG data of all subjects under all conditions are contacinated to form a matrix instead from an individual subject in the step 1.

### 3.5 Statistical analysis

For the yields of the four approaches, we measure the mean amplitudes of two ERPs at five electrodes (FC3, FCz, FC4, C3, Cz, and C4 electrodes). The time window for P2 and N2 is separately 130-190 ms and 190-310 ms. We use two-ways repeated measurement analysis of variance (rm-ANOVA) with valence (extreme, moderate, and neutral) × negative-category (disgusting and fearful) as within-subject factors to compute the statistical results of the measured mean amplitudes for all trials and increased trial numbers, separately. Greenhous-Geisser method is used for correcting the number of degrees of freedom.

Furthermore, the correponded internal consistency of P2/N2 for the four used techniques is also evaluted by measuring the CCs between the averages of smaller trial numbers and the grand averaged N2/P3 (i.e., all trials are included). The Cronbach’s alpha of increased trial numbers is also computed (Unacceptable: *<* 0.05; Poor: 0.5-0.6; Questionable: 0.6-0.7; Acceotable: 0.7-0.8; Good: 0.8-0.9; Excellet: *>* 0.9). In addition, we also compute CCs of topographies between any two different subjects to assess the intrasubject consistency of ERPs. And then, we perform paired t-test on the CCs between M1 and any one of the other approaches to evalute the performance the proposed technique (i.e., M1).

## 4 Results

### 4.1 Internal consistency of P2/N2

We use CCs to examine the relationships between smaller trial averages and the P2/N2 grand average (see Fig. 2, (a) and (b)). We observe the excellent CCs (*>*0.9) after 25 trial are contained in the average of P2 for M1 and 10 trials for the other three approaches. As for N2, the three PCA approaches have higher CCs than M2 and there are no difference among the three PCA strategies. These reveal that relative narrow ERP (i.e., P2) has a significant effect on M1 than others, but N2 has little influence. The evidences clarify that the yields of P2 are highly similar to grand average (all trials) after 25 trials for M1, 10 trials for both M3 and M4, and 12 trial for M2. Similarly, the equivalent measures as the grand average can be obtained after few trials used for N2 when using the four approaches (10, 10, 10, and 14 trials).

**Fig. 2.**
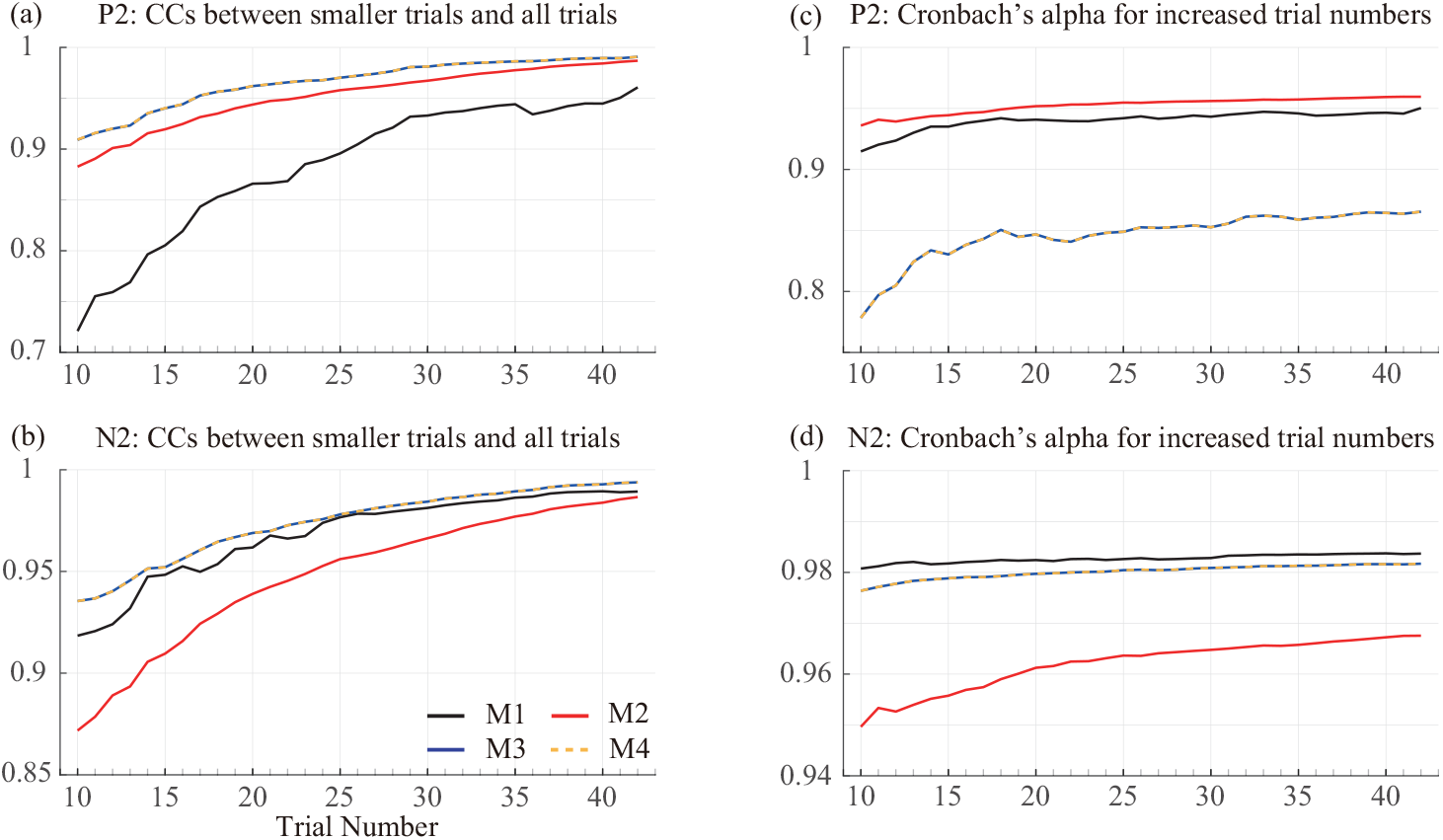
Internal consistency analysis for P2/N2. (a) and (b): Correlation coefficients (CCs) between P2/N2 averages of consecutive increased trials (i.e., 10, 11, …, 41, and 42) and all trials. (c) and (b) are Cronbach’s alpha for adding trials. Electrodes are used for both P2 and N2 at FC3, FCz, FC4, C3, Cz, and C4. M1: Extraction of P2/N2 from single-trial EEG data of an individual subject using temporal-PCA (t-PCA). M2: Conventional time-domain analysis. M3: Group t-PCA for trial-averaged ERP data of all subjects. M4: Group t-PCA for single-trial EEG data of all subjects.

We also compute the Cronbach’s alpha of P2 and N2 for the four approaches, respectively, when trials are added to the averages (see Fig. 2 (c) and (d)). All of them have a growth trend. In order to invistigate excellet Cronbach’s alpha (*>*0.9) for P2, not less than 10 trials should be averaged when using ‘M1’ and ‘M2’. The results of N2 indicate that all used approaches show excellent Cronbach’s alphas for the varied numbers of trials. M1 has highest values than other approaches, especially better than M2.

Noted that both CCs and Cronbach’s alpha of P2/N2 for the two group PCA strategies (i.e., M3 and M4) are completely coincident.

### 4.2 Effect of trial numbers on the PCA factor numbers of P2/N2

Fig. 3 reveals that selected factor numbers may be differ over trials when using M1 to decompose single-trial EEG data of the same individual. Likewise, when the same trial number is used for individuals, the numbes of the extracted factors associated with P2/N2 are also different.

**Fig. 3.**
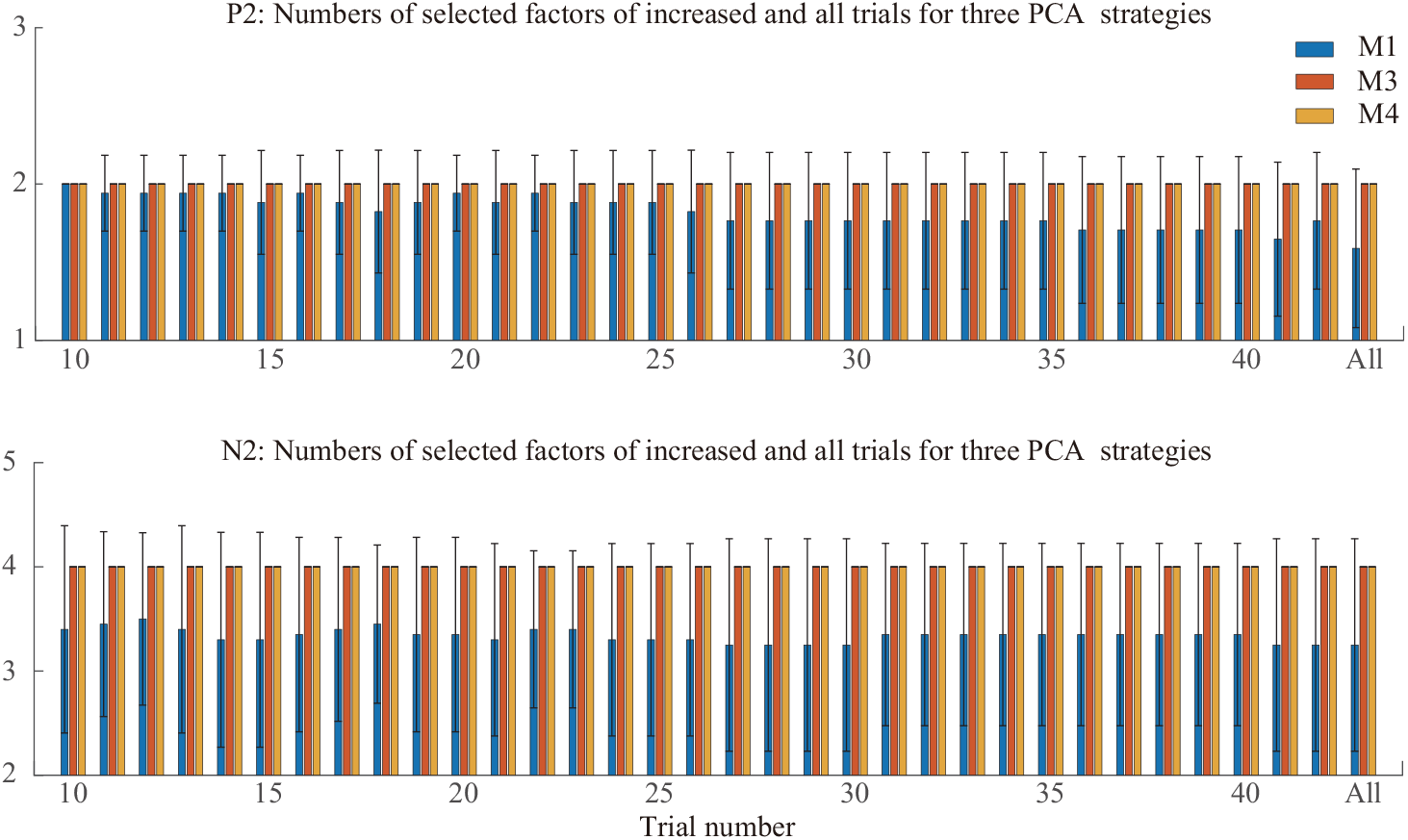
The corresponded numbers of selected factors associated with P2/N2 for the increasing trials and all trials when using three different PCA strategies. M1: Extraction of P2/N2 from single-trial EEG data of an individual subject using temporal-PCA (t-PCA). M3: Group t-PCA for trial-averaged ERP data of all subjects. M4: Group t-PCA for singletrial EEG data of all subjects.

Additionally, the change of trial numbers does not have effect on the number of the selected factors decomposed from waveforms of P2/N2 when using two group PCA strategies. The selected-factors number for P2/N2 is 2/4, and both M3 and M4 extract the same number of factors from P2/N2. By manually checking the decomposed factors for increasing trials and all trial when using both M3 and M4, we observe that the orders of the selected factors related to P2/N2 may be varied across trials.

### 4.3 Trial numbers influence the statistical results of P2/N2

To evaluate the influences of varied trials on statistically analyzed results, we apply two-ways rm-ANOVA to calculate the statistical results based on the mean amplitudes of P2/N2 that are obtained from increased trial numbers and all trials, separately (see Figs. 4 and 5).

**Fig. 4.**
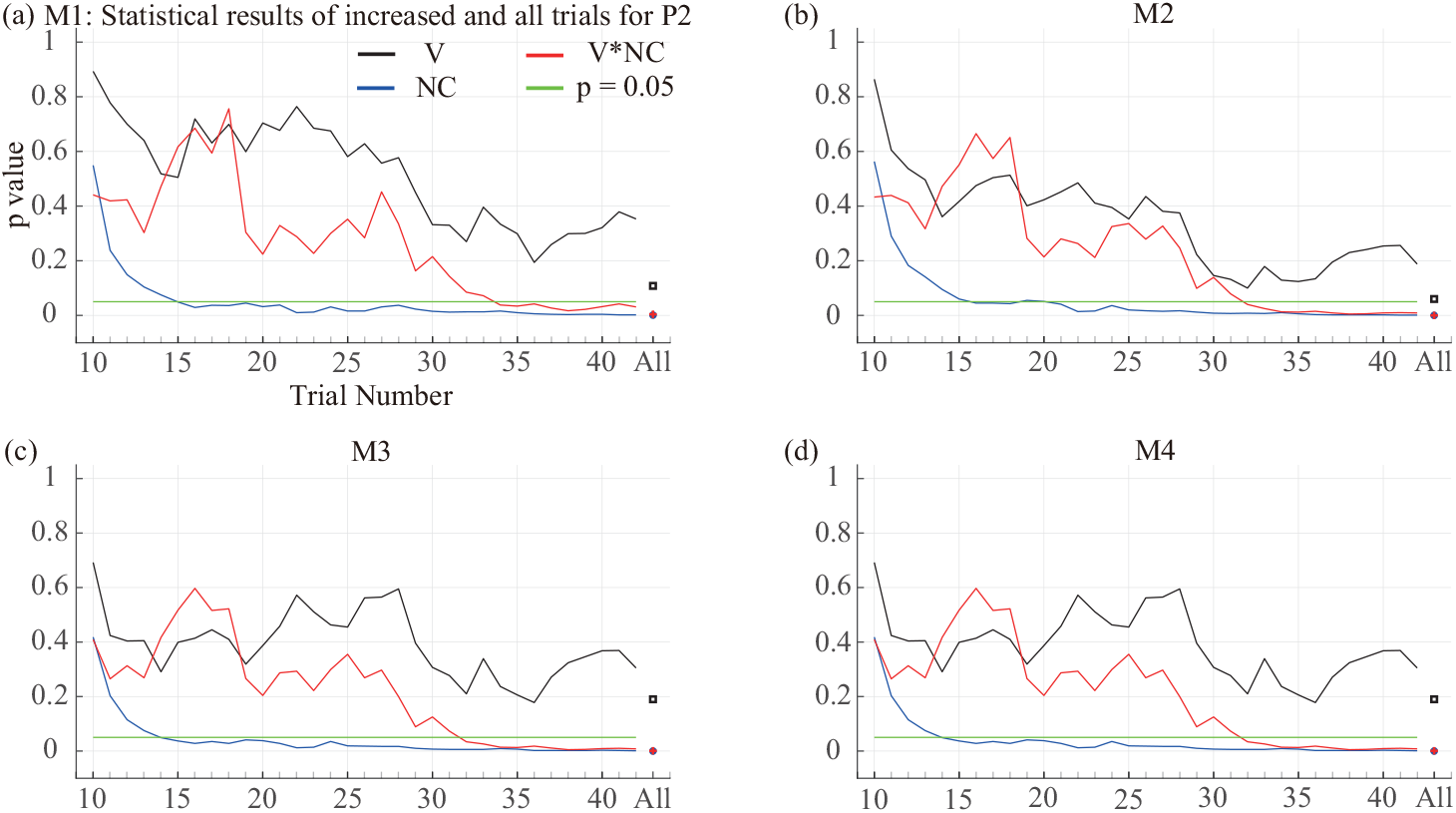
The statistical analysis results for P2 averages of increased trials and grand average (i.e., all trials) when using four different techniques. Valence (V) *×* Negative-category (NC) are the within-subject factors. V∗NC is the interactive effect between valence and negativecategory factors.

**Fig. 5.**
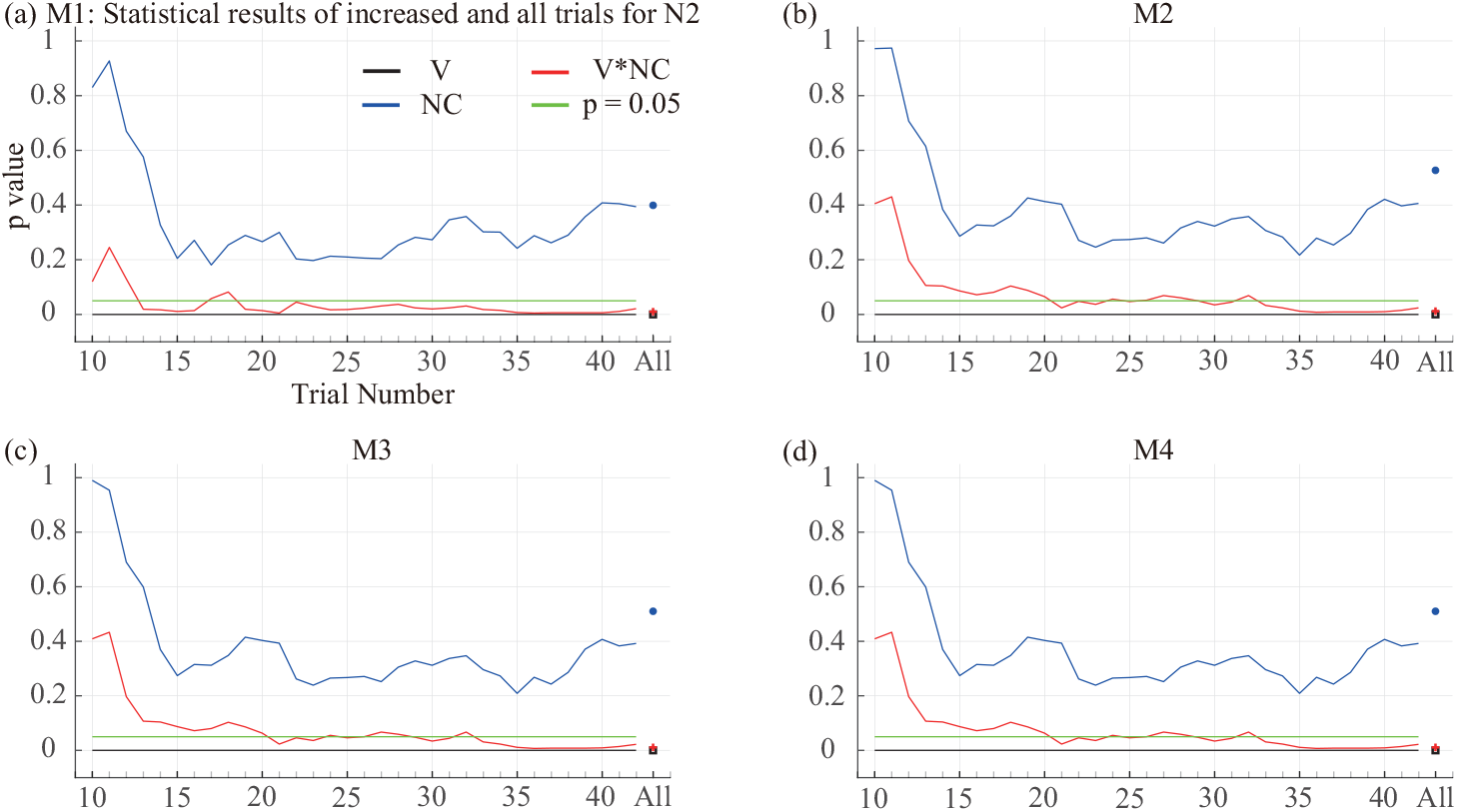
The statistical analysis results for N2 averages of increased trials (from 10 to 42) and grand average (i.e., all trials) when using four different techniques. Valence (V) and Negative-category (NC) are the within-subject factors. V∗NC means the interactive effect between valence and negative-category factos.

Fig. 4 reveals the P2’s statistical results of the four used approaches, we find that the main effects of Valence/Negative-category and their interaction show decreased trends when more trials are added to the average. The changes of all curves become slow down. These trends suggest that adding trials have a little effect on the statistical results after a certain trial number, for example, 35 trials. In other words, we can obtain the equal results to these of all trials after this certain trial number. For the four used techniques, we apply 34 (M1)/32(M2)/32(M3)/32(M4) trials to yield a comparable results as obtained from all trials (see Tab. 1), respectively.

**Table 1.** The statistical results of P2 (130-190ms) for four used techniques when all trials are averaged.

| Methods | *V* |  |  | *NC* |  |  | *V \* NC* |  |  |
| --- | --- | --- | --- | --- | --- | --- | --- | --- | --- |
| | $F_{(2,15)}$ | $\eta_p^2$ | *p* | $F_{(1,16)}$ | $\eta_p^2$ | *p* | $F_{(2,15)}$ | $\eta_p^2$ | *p* |
| *M1* | 2.387 | 0.130 | 0.108 | 17.868 | 0.528 | **0.001** | 7.245 | 0.312 | **0.004** |
| *M2* | 3.079 | 0.161 | 0.060 | 19.392 | 0.548 | **< 0.001** | 10.355 | 0.393 | **< 0.001** |
| *M3* | 1.752 | 0.099 | 0.190 | 21.303 | 0.571 | **< 0.001** | 10.484 | 0.396 | **0.001** |
| *M4* | 1.752 | 0.099 | 0.190 | 21.303 | 0.571 | **< 0.001** | 10.484 | 0.396 | **0.001** |

Similarly, in terms of N2 (Fig. 5), we also observe the downward trends for both main effect of Valence/Negative-category and their interaction when the number of trials is increasing. However, few trials for M1 (19) are used to produce the stable and equative results as all trials (see Tab. 2) for M1 compared with other three techniques (all of them are 33).

**Table 2.** The statistical results of N2 (190-310ms) for four used techniques when all trials are averaged.

| Methods | *V* |  |  | *NC* |  |  | *V \* NC* |  |  |
| --- | --- | --- | --- | --- | --- | --- | --- | --- | --- |
| | $F_{(2,18)}$ | $\eta_p^2$ | *p* | $F_{(1,19)}$ | $\eta_p^2$ | *p* | $F_{(2,18)}$ | $\eta_p^2$ | *p* |
| *M1* | 24.397 | 0.562 | **< 0.001** | 0.746 | 0.038 | 0.399 | 5.553 | 0.226 | **0.010** |
| *M2* | 27.431 | 0.591 | **< 0.001** | 0.415 | 0.021 | 0.527 | 5.081 | 0.211 | **0.012** |
| *M3* | 27.455 | 0.591 | **< 0.001** | 0.452 | 0.023 | 0.510 | 5.198 | 0.215 | **0.011** |
| *M4* | 27.455 | 0.591 | **< 0.001** | 0.452 | 0.023 | 0.510 | 5.198 | 0.215 | **0.011** |

### 4.4 Correlation coefficients of topographies depend on the analyzed ERPs

We also assess the internal subject consistency of P2/N2 by measuring the CCs of topographies (‘spatial CCs’) between any two different subjects (see Figs. 6 (a) and 7 (a)). Meanwhile, paired t-test is implemented to obtain discriminations between M1 and any one of the other three approaches (see Figs. 6 (b) and 7 (b)).

**Fig. 6.**
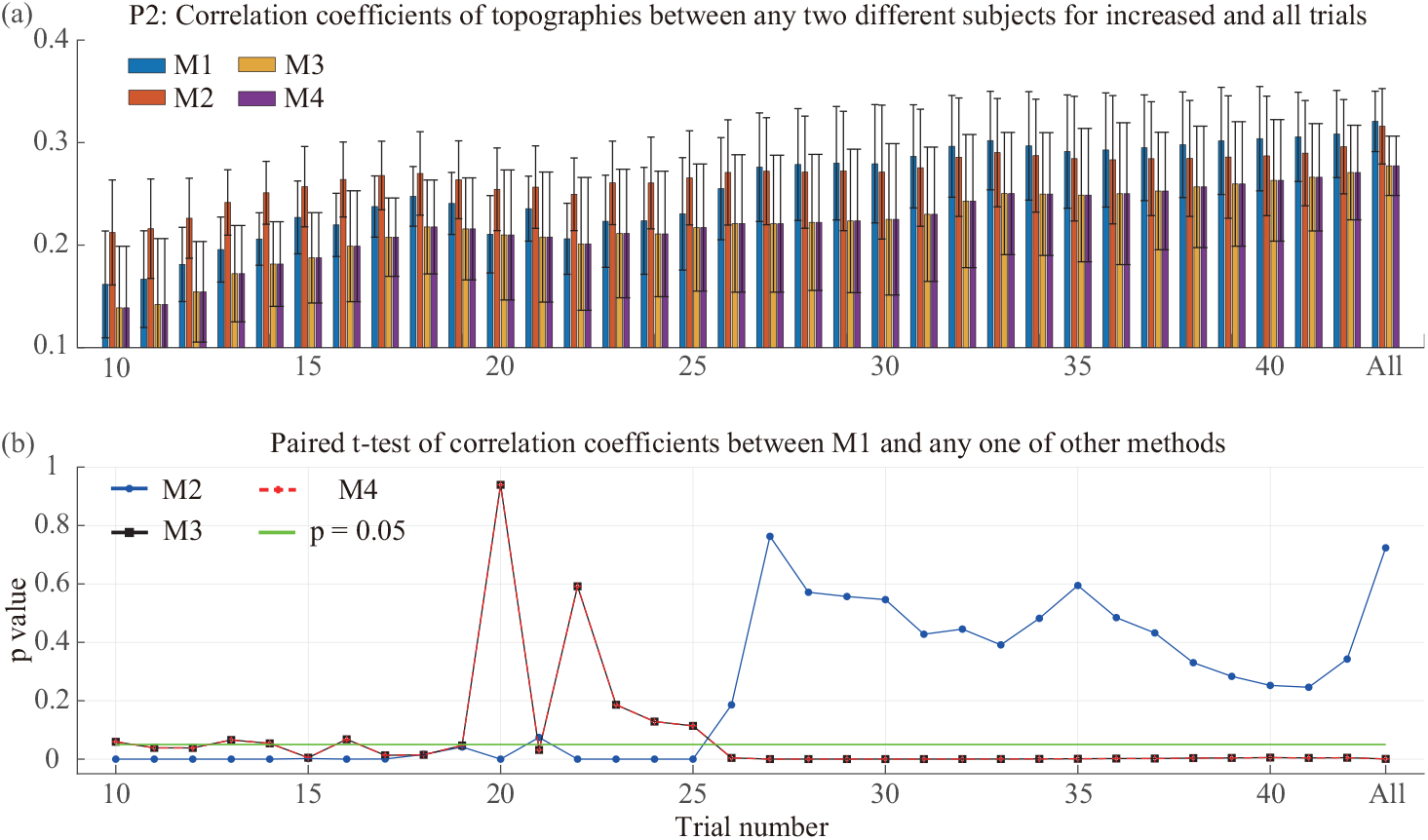
(a) The correlation coefficients (CCs) of P2’s topographies between any two different subjects for adding trial numbers and all trials. (b) Paired t-test of CCs between M1 and any one of the other three methods.

**Fig. 7.**
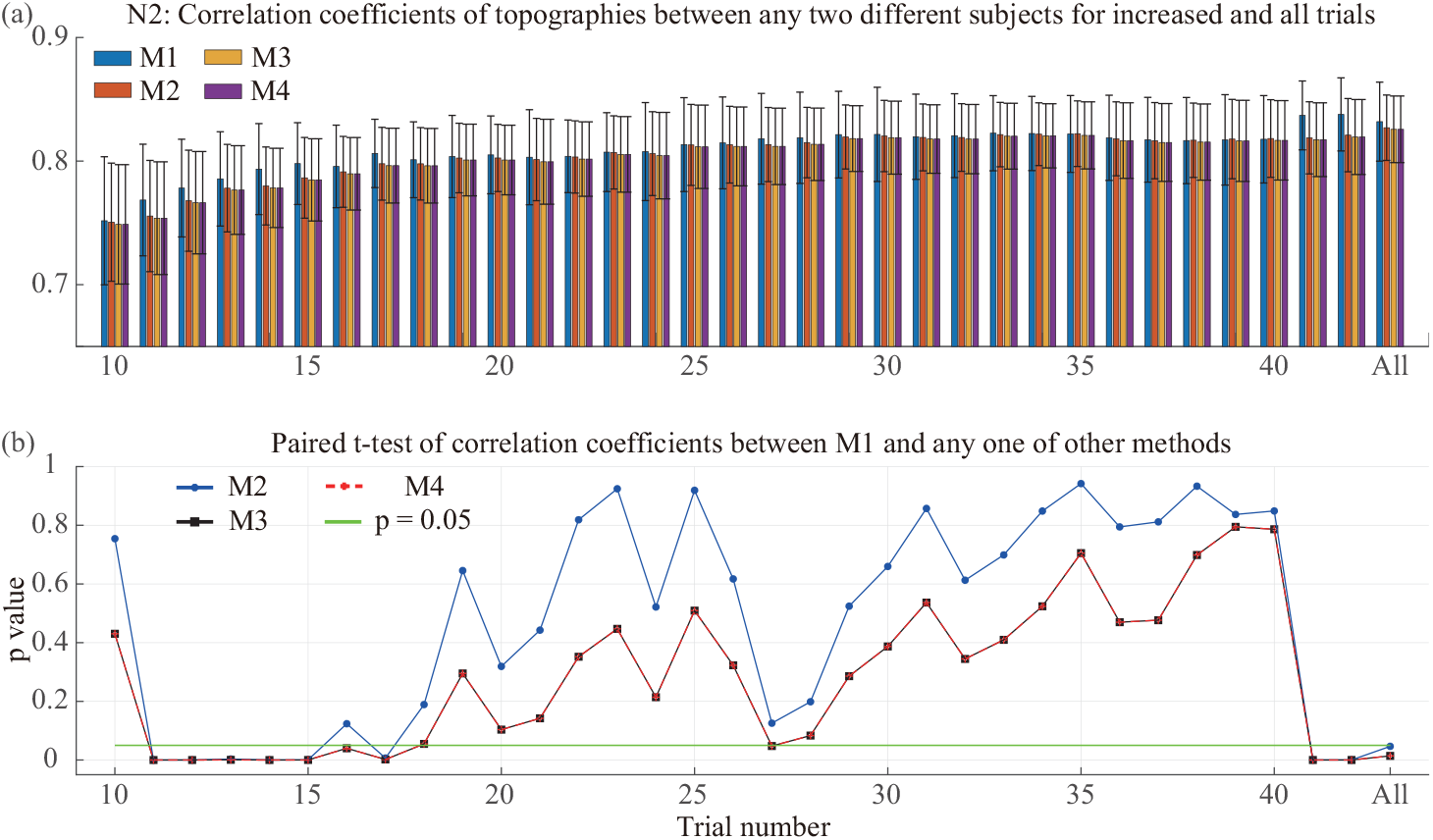
(a) The correlation coefficients (CCs) of N2’s topographies between any two different subjects for increasingly trial numbers and all trials. (b) Paired t-test of CCs between M1 and any one of the other three alternative methods.

For P2 (see Fig. 6), both of M1 and M2 show higher spatial CCs than other two approaches across trials. A significant difference between M1 and M3 or M4 is not observed until more than 26 trials are involved in averages. By contrast, spatial CCs of M2 are significant higher than these of M1 when less than 26 trials are averaged. After 26 trials, although there is no difference between M1 and M2, M1 has excellent values than M2 from 26 trials to 42 trials (also for all trials).

With regard to N2 (see Fig. 7), the mean spatial CC of M1 is higher than that of the other three approaches. The paired t-test results indicate that the spatial CCs of M1 can be significantly improved when 11, 12, 13, 14, 41, 42, and all trials are used compared with other approaches.

Furthermore, the spatial CC of P2 (about 0.25) is lower than N2 (roughly 0.75) for all approaches, suggesting that N2 is more stable than P2 among subjects. In should be also noted that spatial CCs/t-test of P2/N2 for M3 and M4 are no difference across trials.

## 5 Conclusion and discussion

The current study investigates two issues by using a modified oddball emotional experiment with two factors. On the one hand, we examine the influence of trial numbers on the PCA decomposition of P2/N2, which is to obtain stable and reliable ERPs from minimum trial number (i.e., 10, 11, … 41, and 42). On the other hand, we study the discriminations among three PCA strategies/ convenitonal time-domain analysis for the same EEG data. In other words, we employ PCA to extract ERPs of interest from an invidual subject or from all subjects.

In order to realize both purposes, we compute the CCs of P2/N2 between averages with few number of trials and the grand average (i.e., all trials) as used in previous studies (Olvet and Hajcak, 2009; Thigpen et al., 2017; Cohen and Polich, 1997; Rietdijk et al., 2014; Clayson and Larson, 2013). For CC, an excellent yield of P2 can be produced for M2/M3/M4 when minimum trial number is 10, but we only observe similar result for M1 after trial number exceeds 25 trials. By contrast, we obtain the excellent CC of N2 after 10 trials are included in average for three PCA strategies, and 14 trials used for M2.

Also, we calculate the Cronbach’s alpha for increased trials (Olvet and Hajcak, 2009; Kleene et al., 2021; Fischer et al., 2017; Rietdijk et al., 2014; Clayson and Larson, 2013), and obtain the statistical analysis results for different trial numbers and all trials. Regarding Cronbach’s alpha, we obtain excellent alpha values of P2 over all trial numbers for M1/M2 but not for both group PCA strategies (M3 and M4) (see Fig. 2 (c)). As shown in Fig. 2 (d), all alpha values of N2 for the four techniques reach excellent level when more than 10 trials are used. In terms of the statistical results, Fig. 4 reveals that stable and reproductive results of P2 for four used techniques can be obtained after about 35 trials are applied to the average procedure. However, only 19 trials can be obtained the considerable result of N2 as all trials when using M1, and approximately 33 trials for the other three techniques (see Fig. 5).

Additionally, the CCs of topographies (we following call it spatial CCs) among different subjects are also used to assess the internal-subject consistent of P2/N2 of interest and those are also used to measure the performances of different techniques. In terms of an ERP, it is characterized by both a specific time course and a specific topography. For example, in a typical oddball paradigm, P3a can be elicited about 250-280ms at frontal-central sites (Polich, 2007; Comerchero and Polich, 1999). Generally, in an ERP experiment, ERPs are often quantified from tens or hundreds of subjects, and thus, we expect that brain activities related to stimulus onset can be observed from all subjects so that the related experiment is reproducible and reliable. A reliable ERP response means that higher CCs of topographies between any two different subjects can be obtained. As shown in Fig. 6 (a) and (b), regarding P2, the spatial CCs for the proposed technique are significantly higher than those of the other two group strategies (M3 and M4) after the number of trials exceeds 25 but not for M2. Likewise, for N2, although there is no difference among four techniques in most of trials, the spatial CC values of the proposed technique are higher than others (see Fig. 7).

The main difference between the proposed technique (M1) and the conventional time-domain analysis (M2) is that M1 explores both temporal and spatial characteristics of P2/N2, but M2 only uses the temporal (i.e., amplitude) property of P2/N2 and the obtained results are still mixtures. Moreover, compared with the other two group PCA strategies (M3 and M4) in which ERPs of interest are extracted from all subjects, the proposed technique explores the desired ERPs from the single-trial EEG of each subject. Traditional group PCA strategies (M3 and M4) assume that the numbers and the orders of sources are the same for all subjects (Dien, 2012), but M1 allows both parameters to be varied among subjects (see Fig. 3). Noticeably, the core ideas of M3 and M4 are the same and PCA is a linear decomposition, and thus, we believe that both PCA strategies are no difference when using the same estimated source number. This hypothesis is also confirmed by the results of the CC, Cronbach’s alpha, statistical analysis, and spatial CC (see Figs. 2, 4, 5, 6, and 7).

In summary, the results in the current study lead to the following four suggestions: (1) ERPs of interest can be efficiently extracted from single subject by PCA decomposition rather than from the datasets of all subjects simultaneously (which is often used in PCA Toolkit (Dien, 2010a)). (2) The ERP component that have higher spatial CC use few trial number to yield the highly similar results to all trials compared with other ERPs with lower spatial CC. For instance, 19 trials can obtain an internally consistent measure for N2 and 35 trials for P2 when using the proposed apporach. (3) ERP components that have a wider time window are easily to be decomposed into more factors by PCA than other that have a narrow time range. For example, N2 is decomposed into four factors by M3/M4 and P2 is decomposed into two factors (see Fig. 3). (4) The yields of M3 which extracts ERPs from averaged ERP data of all subjects are the same to the M4 that explores ERPs from single-trial EEG at the group level. (5) The similarities of topographies among different subjects can be used as both a criterion to evaluate the performance of the used techniques and a method to judge whether the analysed ERPs are repeatable and reliable or not.

Furthermore, there are some potential limitations to the present study. Firstly, we only perform PCA on the spatial-stacked matrix (i.e., temporal-PCA) to extract P2/N2 of interest in the current study. Although the proposed technique produces much better overall results than those of all the other techniques, this conclusion is not further validated in the applications of spatial PCA. Secondly, the comparison of yields between PCA and ICA is not included in this study. Many applications of ICA indicated that ERPs of interest can also be efficiently extracted from both single-trial EEG and averaged ERP datasets (Lee et al., 2016; Wessel, 2018; Cong et al., 2013; Rissling et al., 2014). It is worth investigating the information of the desired ERP using ICA from either average traces or single-trial traces. Thirdly, the identification of PCA-extracted factors associated with the desired ERP seems to be a subjective way in this study, which is only taken its temporal and spatial properties into consideration. As described in application of ICA on the single-trial EEG of single subject (Rissling et al., 2014), we can also identify the PCA-extracted factors related to ERPs of interest by using characteristics of the factor topography, factor waveform, factor spectra, and factor dipoles. Lastly, we merely invistigate the effects of trial numbers on the internal consistency of ERPs and statistical results, the interactive effects between subjects and trials are not further studied as in past report (Boudewyn et al., 2018) because few subjects (i.e., 22) are invovled in current study.

## Acknowledgments

This work was supported by National Natural Science Foundation of China (Grant No.91748105), National Foundation in China (No. JCKY2019110B009 & 2020-JCJQ-JJ-252) and the Fundamental Research Funds for the Central Universities [DUT20LAB303 & DUT20LAB308] in Dalian University of Technology in China, and the scholarships from China Scholarship Council (No.201806060165). This study is to memorize Prof. Tapani Ristaniemi for his great help to Guanghui Zhang, Fengyu Cong and the other authors.

## Footnotes

1 factor is used here to represent PCA-extracted component by PCA + rotation method, which is to avoid for confusion with ERP component.

